## Appendix 1: Search Criteria for "Do organisms need an impact factor? Citations of key biological resources including model organisms reveal usage patterns and impact"

**AGSC Search Criteria:**

<https://scholar.google.com/scholar?start=110&q=%22RRID:SCR_006372%22+OR+%22SCR_006372%22+OR+%22nlx_152125%22+OR+%22www.ambystoma.org/genetic-stock-center%22+OR+%22orip.nih.gov/comparative-medicine/programs/vertebrate-models%22+OR+%22RRID:AGSC%22+OR+%22AGSC_%22++OR+%22AGSC+Cat%22+OR+%22AGSC+catalog%22+OR+%22Ambystoma+Genetic+Stock+Center%22%22&hl=en&as_sdt=0,5&as_ylo=2012&as_yhi=2022&as_vis=1>

**NXR Search Criteria:**

<https://scholar.google.com/scholar?hl=en&as_sdt=0%2C5&as_ylo=2011&as_yhi=2022&q=%22NXR+frog%22+OR+%22RRID%3ASCR_013731%22+OR+%22SCR_013731%22+OR+%22www.mbl.edu%2Fxenopus%2F%22+OR+%22orip.nih.gov%2Fcomparative-medicine%2Fprograms%2Fvertebrate-models%22+OR+%22National+Xenopus+Resource%22+OR+%22RRID%3ANXR_%22+OR+%22RRID%3ANXR%22+OR+%22NXR_%22+&btnG=>

**ZIRC Search Criteria:**

<https://scholar.google.com/scholar?start=440&q=%22RRID:SCR_005065%22+OR+%22SCR_005065%22+OR+%22zebrafish.org%22+OR+%22zebrafish.org/home/guide.php%22+OR+%22RRID:ZIRC%22+OR+%22ZIRC+Cat%22+OR+%22ZIRC+catalog%22+OR+%22ZIRC_%22&hl=en&as_sdt=0,5&as_ylo=2011&as_yhi=2022&as_vis=1>

**MMRRC Search Criteria:**

<https://www.ncbi.nlm.nih.gov/pmc/?term=%22RRID%3ASCR_016448%22+OR+%22SCR_016448%22+OR+%22mmrrc.ucdavis.edu%2F%22+OR+%22MMRRC%22+OR+%22RRID%3AMMRRC+UCD%22+OR+%22Mutant+Mouse+Resource+and+Research+Center%22+OR+%22Mutant+Mouse+Resource+%26+Research+Center%22>
